# RANKOR: Direct Drug Prioritization from Bulk and Single-Cell Transcriptomic Signatures

**DOI:** 10.64898/2026.05.20.726471

**Authors:** Nikoletta Katsaouni, Marcel H. Schulz

**Author notes:** Contributing authors.

## Abstract

**Background:** Prioritizing therapeutics from transcriptomic data remains a key challenge in precision medicine. Signature reversal approaches, most commonly implemented through Gene Set Enrichment Analysis (GSEA), have been widely used to match disease signatures to candidate drugs. However, enrichment-based methods can be sensitive to noise and are restricted to previously profiled compounds

**Methods:** We developed RANKOR, a machine-learning framework designed to rank candidate drugs directly from transcriptomic signatures. Rather than predicting full expression profiles, RANKOR learns structured latent representations of transcriptional responses alongside chemical structure, enabling prioritization from standardized signatures derived from disease states or treatment perturbations. The framework is applicable to both bulk and single-cell transcriptomic data.

**Results:** Across large-scale perturbational datasets, RANKOR achieved consistently lower median ranks than similarity- and distance-based approaches, while showing performance comparable to, and in some settings improved over, GSEA. The model generalized across unseen cell types and retained performance in single-cell settings, where it provided more consistent prioritization than existing approaches, such as ASGARD. RANKOR further enabled prioritization of transcriptionally unseen compounds through chemical-space embedding and achieved substantially reduced computation times. Robustness analyses demonstrated stable performance under moderate noise and degradation under extreme perturbation or gene shuffling. Gene attribution analyses indicated that prioritization decisions are driven by coherent and mechanism-relevant transcriptional programs.

**Conclusions:** RANKOR provides a scalable framework for transcriptomics-guided drug prioritization that can complement and extend existing approaches, such as GSEA. It can also support therapeutic hypothesis generation from bulk and single-cell data while leveraging the generalisability and computational efficiency of machine learning models.

## 1 Introduction

Precision medicine aims to select therapies that modulate the molecular processes driving disease in individual patients or clinically meaningful patient subgroups (Strianese et al. 2020; Wang and Wang 2023). In many settings—including cancer, immune disorders, and rare disorders—disease biology is reflected not by single genetic alterations but by coordinated changes in transcriptional programs and pathway activity (Johansson et al. 2023). Transcriptomic profiling therefore provides a powerful intermediate phenotype, integrating genetic, epigenetic, and environmental influences into a measurable cellular state that can be linked to therapeutic hypotheses (Santos et al. 2017; Pushpakom et al. 2019; Trapotsi et al. 2022). Recent single-cell and single-nucleus transcriptomic studies in patient-derived samples have demonstrated how cellular heterogeneity and disease-specific transcriptional states can directly inform therapeutic stratification, including in hematological malignancies and autoimmune disease (Biradar et al. 2025; Huang et al. 2026).These works highlight the growing need for drug prioritization methods that can operate directly on patient-level bulk and single-cell transcriptomic signatures, rather than relying on predefined gene sets, which may overlook patient-specific transcriptional heterogeneity, or on matched reference drug profiles, which are not available for all compounds and experimental conditions.

Clinically, this reflects a common scenario: a patient or molecularly defined disease subgroup exhibits a transcriptomic signature, and the goal is to identify candidate therapies—including compounds not directly profiled in a matched experimental setting—for experimental validation or translational follow-up. Once such a molecular profile is obtained, a central translational question arises: which therapies are most likely to reverse the dysregulated biological programs observed in that patient or disease subgroups. This prioritization task is highly consequential yet experimentally intractable. Systematically screening large drug libraries across diverse patient-specific contexts is infeasible. Consequently, computational strategies that connect disease-associated gene expression signatures to therapeutic hypotheses have become an important component of modern repurposing and discovery pipelines (Chen et al. 2026; Pushpakom et al. 2019; Cousins et al. 2024; Musa et al. 2018).

Recent years have seen rapid expansion of machine learning applications leveraging large-scale perturbational and chemical datasets. Predictive models have been developed to infer drug mechanisms of action, predict transcriptional responses following compound exposure, and estimate potency or sensitivity metrics from gene expression and structural features (Yu and Fan 2025; Yu et al. 2025; Jiang et al. 2024, 2022; Gao et al. 2021; Jang et al. 2021). Representation learning frameworks further aim to embed drugs and transcriptional states into shared latent spaces that capture pharmacological similarity and regulatory relationships. Additionally, recent integrative machine learning frameworks combining perturbational atlases with single-cell transcriptomics further demonstrate how drug–cell state relationships can be modeled at high resolution (Gupta et al. 2025).

While these advances provide powerful tools for modeling drug-induced transcriptional biology, they do not directly solve the translational prioritization problem. Many predictive frameworks output high-dimensional gene expression reconstructions, similarity scores, or abstract potency estimates that require additional interpretation before therapeutic hypotheses can be formulated (Qi et al. 2024; McFarland et al. 2020; Li et al. 2021; Hetzel et al. 2022; Lotfollahi et al. 2023; Piran et al. 2024; Szalata et al. 2024). Moreover, predictive accuracy does not necessarily translate into reversal of disease-associated molecular programs. Several additional limitations constrain clinical applicability. Predictions depend on training coverage and may not generalize to unseen compounds or biological contexts. Model outputs can lack mechanistic interpretability, and predicted transcriptional effects do not inherently encode therapeutic directionality. Thus, despite methodological sophistication, many machine learning systems address adjacent predictive problems rather than the direct task of ranking therapies capable of reversing disease states.

However, a subset of transcriptomics-driven drug repurposing methods relies on the principle of *signature reversal* (Koudijs et al. 2019; Chen et al. 2017; Shukla et al. 2021; Chalkioti et al. 2025; Ahmed et al. 2022; O’Donovan et al. 2021). Disease states can be characterized by coordinated up- and down-regulation of genes relative to healthy tissue, while drug perturbations induce their own transcriptional programs. If a compound drives expression changes in the opposite direction of a disease signature, it may counteract disease-associated molecular mechanisms.

This concept was formalized by the Connectivity Map, which demonstrated that similarity and anti-correlation between disease and drug signatures can recover known therapeutic relationships and suggest repurposing opportunities (Lamb et al. 2006; Subramanian et al. 2017). Connectivity mapping has since been applied across therapeutic areas and biological modalities. Early demonstrations integrated public disease expression compendia with perturbational signatures to predict candidate indications (Sirota et al. 2011), and subsequent studies validated computational repositioning predictions in preclinical systems such as inflammatory bowel disease (Dudley et al. 2011).

The expansion of perturbational resources, particularly under the LINCS program and the L1000 platform, enabled systematic large-scale application of connectivity principles (Subramanian et al. 2017). Community platforms such as iLINCS (Pilarczyk et al. 2022) and L1000CDS2 (Duan et al. 2016) operationalize signature search workflows to identify compounds predicted to mimic or reverse query profiles. Connectivity-style analyses have also been used to organize drug perturbations into pharmacological networks and infer mechanisms of action (Iorio et al. 2010; Carrella et al. 2014).

Among signature-based methods, Gene Set Enrichment Analysis (GSEA) is widely used to interpret differential expression through pathway-level gene sets and has become a standard analytic component in translational transcriptomics (Subramanian et al. 2005). Enrichment-based formulations have also been adapted for mechanism-centric therapeutic prioritization and drug grouping (Garana et al. 2023).

Despite interpretability and broad adoption, enrichment- and correlation-based signature reversal approaches face important limitations. First, they rely heavily on gene ranking or thresholding, making outputs sensitive to noise and instability in differential expression estimates—a challenge amplified in heterogeneous cohorts and single-cell settings (Cousins et al. 2024; Tham et al. 2022). Second, many formulations treat genes as largely independent units, underrepresenting gene–gene dependencies and nonlinear regulatory effects that shape pathway activity and drug response (Trapotsi et al. 2022; Del Real and Rubio 2023). Third, classical connectivity analysis depends on the availability of reference transcriptional profiles: compounds which are not experimentally profiled in relevant contexts cannot be directly evaluated (Subramanian et al. 2017; Duan et al. 2016).

Taken together, these observations highlight a conceptual gap between predictive modeling and translational prioritization. Machine learning frameworks model transcriptional or pharmacological effects (Jiang et al. 2022, 2024; Yu and Fan 2025), but do not directly rank therapeutics for reversal of disease states. Signature-based approaches generate ranked hypotheses but remain constrained by noise sensitivity and experimental coverage.

Here we introduce **RANKOR**, a framework designed to treat drug prioritization itself as the primary objective. Given a transcriptomic signature derived from disease–healthy comparisons, treatment perturbations, or patient single-cell profiles, RANKOR directly ranks candidate compounds according to their alignment with a desired biological shift. By learning structured and aligned representations of transcriptional responses and chemical structure, the framework enables prioritization of both profiled and previously unseen drugs.

In the remainder of this study, we evaluate RANKOR across large-scale perturbational datasets and patient-derived transcriptomic contexts, assessing prioritization performance, computational efficiency, robustness to noise, and biological interpretability. The source code of RANKOR for data processing, analysis and direct drug prioritation is publicly available at: [RANKOR](https://github.com/SchulzLab/RANKOR)

## 2 Results

### 2.1 RANKOR enables direct drug prioritization from bulk and single-cell transcriptomic signatures

We developed RANKOR, a representation learning–based framework for transcriptomics-driven drug prioritization. RANKOR connects gene expression signatures to therapeutic candidates by embedding biological and chemical information into aligned latent spaces, enabling direct and efficient drug ranking from transcriptomic data The framework is built to connect transcriptomic signatures to therapeutic candidates through structured representations of biological and chemical space, enabling direct drug ranking from gene expression data. To provide an overview of the RANKOR framework and its application to drug prioritization, Fig. 1 summarizes both the learning phase and the downstream ranking workflow.

**Fig. 1.**
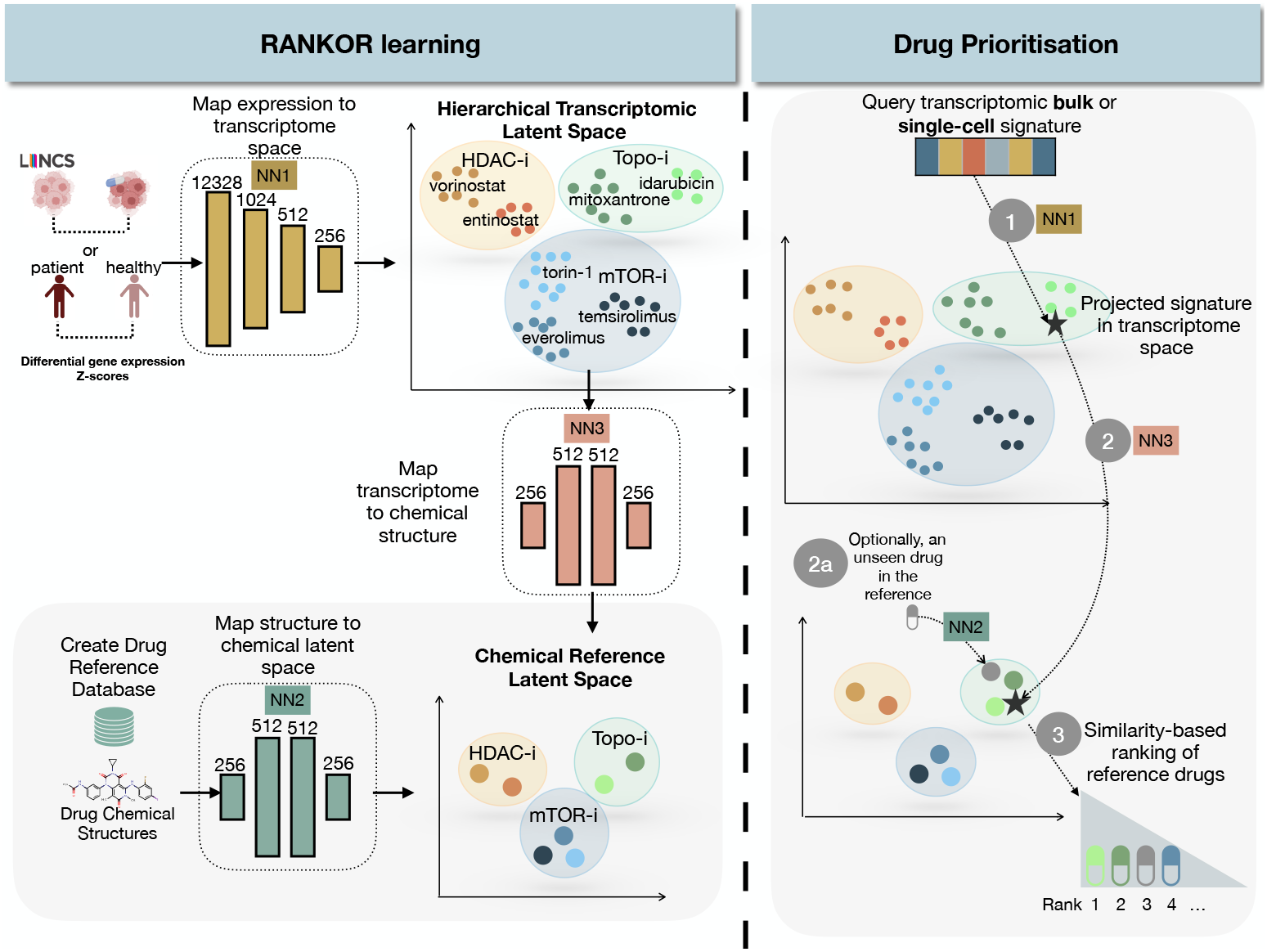
RANKOR framework for transcriptomics-driven drug prioritization. **Left:** During training, gene expression signatures are organized into a hierarchical transcriptomic latent space structured by shared mechanisms of action, while drug chemical structures are embedded into a complementary chemical latent space with one representation per compound. A mapping between the two spaces links transcriptional responses to chemical profiles. NN1 (transcript encoder) learns a representation of gene expression signatures that reflects biological mechanisms and perturbation similarity. NN2 (chemical encoder) transforms molecular features into a compact embedding representing each drug. NN3 (mapping network) projects the transcriptomic representation into the chemical space (or vice versa), allowing direct comparison and ranking of candidate drugs based on similarity. **Right:** For drug prioritization, a bulk or single-cell transcriptomic query is projected through the learned spaces and drugs are ranked by similarity in chemical space, yielding an ordered list of candidate compounds, including (optionally) previously unseen drugs.

During the learning phase (Fig. 1, left), RANKOR constructs two complementary latent spaces, representing transcriptomic and chemical information. Integrating these modalities has been shown to improve performance in drug repurposing by capturing both biological response and molecular structure (Stokes et al. 2020; Zhavoronkov et al. 2019).

Differential gene expression signatures—derived either from drug perturbations in reference datasets or from comparisons between disease and healthy states—are first mapped into a hierarchical transcriptomic latent space with a standard 2 layer fully connected neuronal network (NN1). In this space, transcriptional responses organize according to shared mechanisms of action, while preserving biologically meaningful variability across individual compounds, doses, time points, and cellular contexts (Katsaouni and Schulz 2026). Drugs with similar biological effects form coherent regions, whereas individual compounds appear as structured subclusters within these regions. In parallel, chemical structures of reference drugs are embedded into a chemical latent space with another two layer fully connected neural network (NN2), where each compound is represented by a single point. This representation provides a compact, one-vector-per-drug view that preserves mechanistic relationships between compounds. The transcriptomic and chemical latent spaces are aligned through a learned mapping with a third neural network (NN3), allowing multiple transcriptional profiles associated with the same compound to converge toward a unified chemical representation.

Once the latent spaces are established, RANKOR can be applied to drug prioritization tasks (Fig. 1, right). A query transcriptomic signature—originating from bulk or single-cell data—is projected into the transcriptomic latent space and subsequently mapped into the chemical latent space. This projection represents the chemical profile most consistent with the observed transcriptional changes. Candidate drugs are then ranked based on their similarity to the projected query in chemical space, yielding an ordered list of therapeutic candidates.

Importantly, this formulation enables drug prioritization even when the queried drug or condition has not been previously profiled in the reference dataset. By operating through shared latent representations rather than direct signature matching, RANKOR supports generalization to unseen drugs and cellular contexts. The resulting output is a ranked list of candidate compounds, rather than predicted gene expression profiles or enrichment scores, directly aligning with the objective of therapeutic prioritization.

### 2.2 Role of the chemical latent space in RANKOR

The learned chemical latent space plays a central role in RANKOR. By mapping each compound to a single embedding, it provides a common target space to which multiple transcriptomic signatures of the same drug can be aligned during cross-space mapping. Moreover, because chemical embeddings are derived independently of transcriptional data, the same encoder can be applied to compounds that have not been experimentally profiled, enabling their inclusion in the drug prioritization task. Drug ranking is ultimately performed by comparing projected transcriptomic queries to chemical embeddings in this space using similarity measures, as described below.

As illustrated in Fig. 2A, the learned chemical latent space exhibits a structured organization that reflects pharmacological mechanisms of action (MoAs). Compounds that share the same MoA are enforced to cluster together in this space, despite differences in detailed chemical structure. For example, the BCR–ABL inhibitors ponatinib and nilotinib occupy adjacent regions, consistent with their shared kinase-targeting activity. Similarly, the topoisomerase inhibitors pirarubicin and mitoxantrone are positioned in close proximity, forming a distinct neighborhood associated with DNA-interacting agents. Phosphodiesterase inhibitors such as sildenafil and tadalafil also cluster together, despite structural variations between them. These patterns indicate that the learned embeddings capture functional pharmacological similarity rather than superficial structural resemblance alone. In other words, the chemical latent space is organized according to shared biological activity, enabling drugs with common mechanisms—but potentially diverse chemical scaffolds—to be represented in neighboring regions. This structured chemical reference space provides the basis for cross-modal alignment with transcriptomic representations and downstream drug prioritization.

**Fig. 2.**
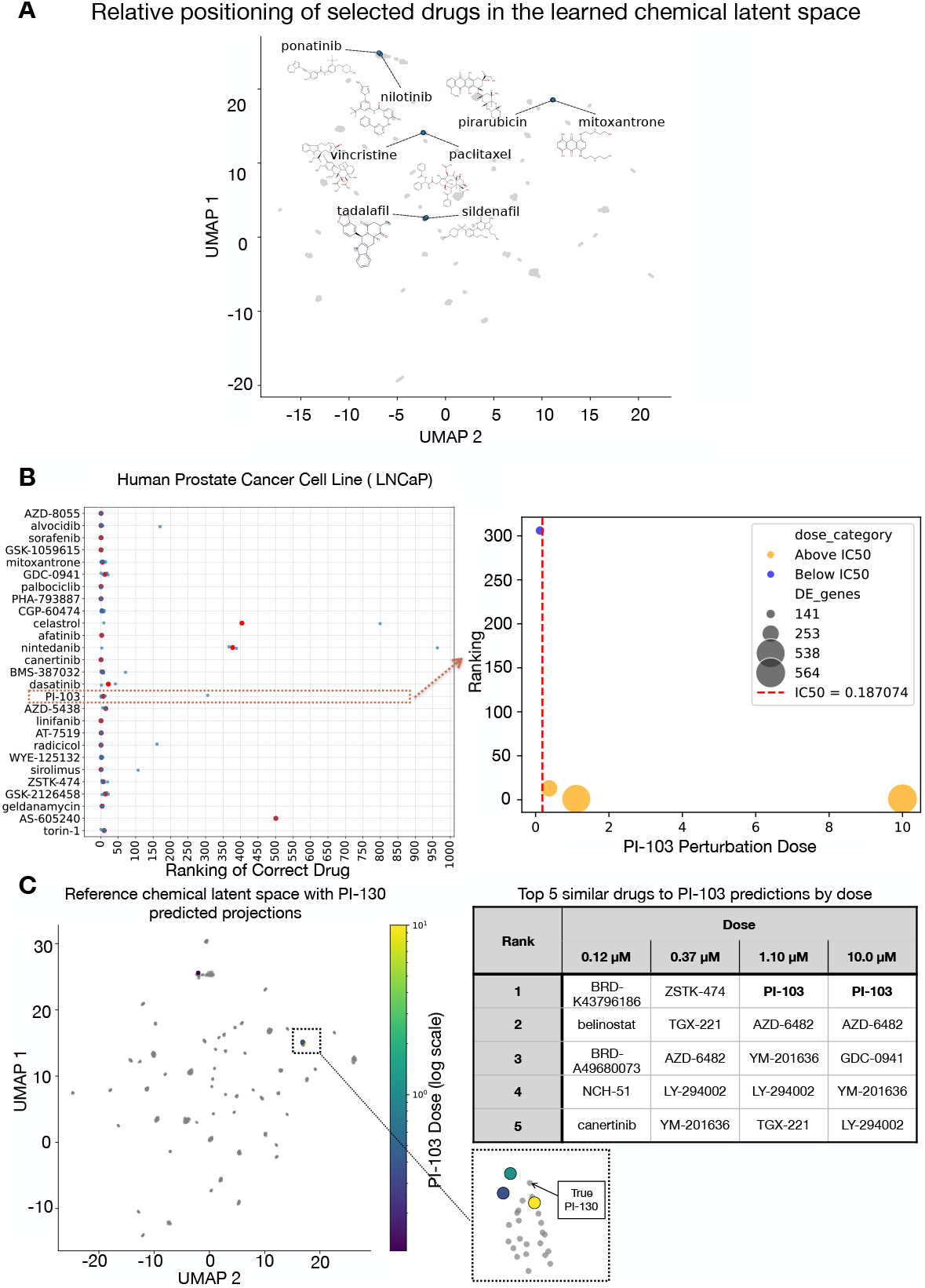
Organization of the chemical latent space learned by RANKOR. **(A)** Two-dimensional visualization of compound embeddings shows clustering by shared mechanisms of action. Kinase inhibitors, topoisomerase inhibitors, phosphodiesterase inhibitors, and microtubule-targeting agents form coherent groups. Compounds with similar pharmacological activity co-localize despite structural differences, indicating that the embedding captures functional rather than purely structural similarity. **(B)** Drug-specific ranking variability in LNCaP cells and dose–response behavior for PI-103. Improved (lower) rankings correspond to stronger transcriptional signal, with limited performance at sub-IC_50_ concentrations as it is shown in the right panel where the x axis corresponds to the applied drug dosage and the y axis to the ranking of the correct drug. **(C)** Projection of PI-103 test perturbations into the chemical latent space across doses, showing convergence toward the true compound embedding with increasing perturbation strength (zoom in on the bottom right). The table on the right summarizes top-ranked candidate drugs at each dose.

### 2.3 Dose-dependent variability reflects biological signal strength

For specific compounds within individual cell lines ranking variability was observed. To investigate this behavior and to fully understand the rationality behind RANKOR prioritisations, we focused on one cell type, the LNCaP prostate cancer cell line (Fig. 2B). For some drugs, correct rankings were consistently low, whereas for others, performance varied substantially depending on perturbation conditions.

Using PI-103 as a representative example (Fig. 2B, right), we observed a clear relationship between perturbation dose, transcriptional signal strength, and ranking performance. At very low concentrations (e.g., 0.12 *µ*M), few genes exhibited differential expression, and the correct compound was poorly ranked. As dose increased, the number of responsive genes grew, and the ranking of the true drug improved markedly. The vertical dashed line indicates the IC_50_ (half-maximal inhibitory concentration), defined as the concentration required to achieve 50% inhibition of biological activity. Doses substantially below the IC_50_ typically induce weak or nonspecific transcriptional changes, limiting recoverable mechanistic signal. As perturbation strength approaches or exceeds IC_50_, the induced transcriptional program becomes more distinct, enabling improved prioritization. Thus, performance heterogeneity may often reflect biological signal availability rather than model instability.

#### Dose-dependent convergence in chemical latent space

To further interpret these findings, we examined the projections of PI-103 perturbations at different doses into the chemical latent space (Fig. 2C). The chemical space is structured by mechanism of action, with each compound represented by a fixed embedding derived independently of transcriptional data. Query signatures are projected into this space and ranked via cosine similarity.

At the lowest dose, the projected embedding was positioned far from the true PI-103 reference embedding. With increasing dose (0.37 *µ*M, 1.10 *µ*M, 10 *µ*M), the projected points progressively moved closer to the true compound location. At the highest dose, the predicted embedding closely overlapped the PI-103 reference.

The top-ranked compounds at each dose further support this interpretation. At low dose, other kinase inhibitors (e.g., ZSTK-474, TGX-221, AZD-6482, GDC-0941) were prioritized. These compounds are known PI3K-pathway inhibitors with overlapping pharmacological activity (Vanhaesebroeck et al. 2012; Kong and Yamori 2010; Raynaud et al. 2009). As dose increased, PI-103 itself emerged as the top-ranked candidate. Thus, even when exact recovery was imperfect under weak signal conditions, RANKOR prioritized pharmacologically coherent alternatives.

### 2.4 RANKOR prioritizes correct compounds across unseen cellular contexts

In order to assess the ability of RANKOR to generalize across biological contexts, we evaluated the framework on unseen cell types. Figure 3A shows the rank of the correct compound across four held-out cell lines (NCIH508, A549, MDA-MB-231, and LNCaP), where lower values indicate better prioritization. RANKOR was compared against enrichment-based signature reversal (GSEA), correlation-based approaches (Pearson and cosine similarity), distance-based matching (Euclidean distance), and set-overlap similarity (Jaccard index).

**Fig. 3.**
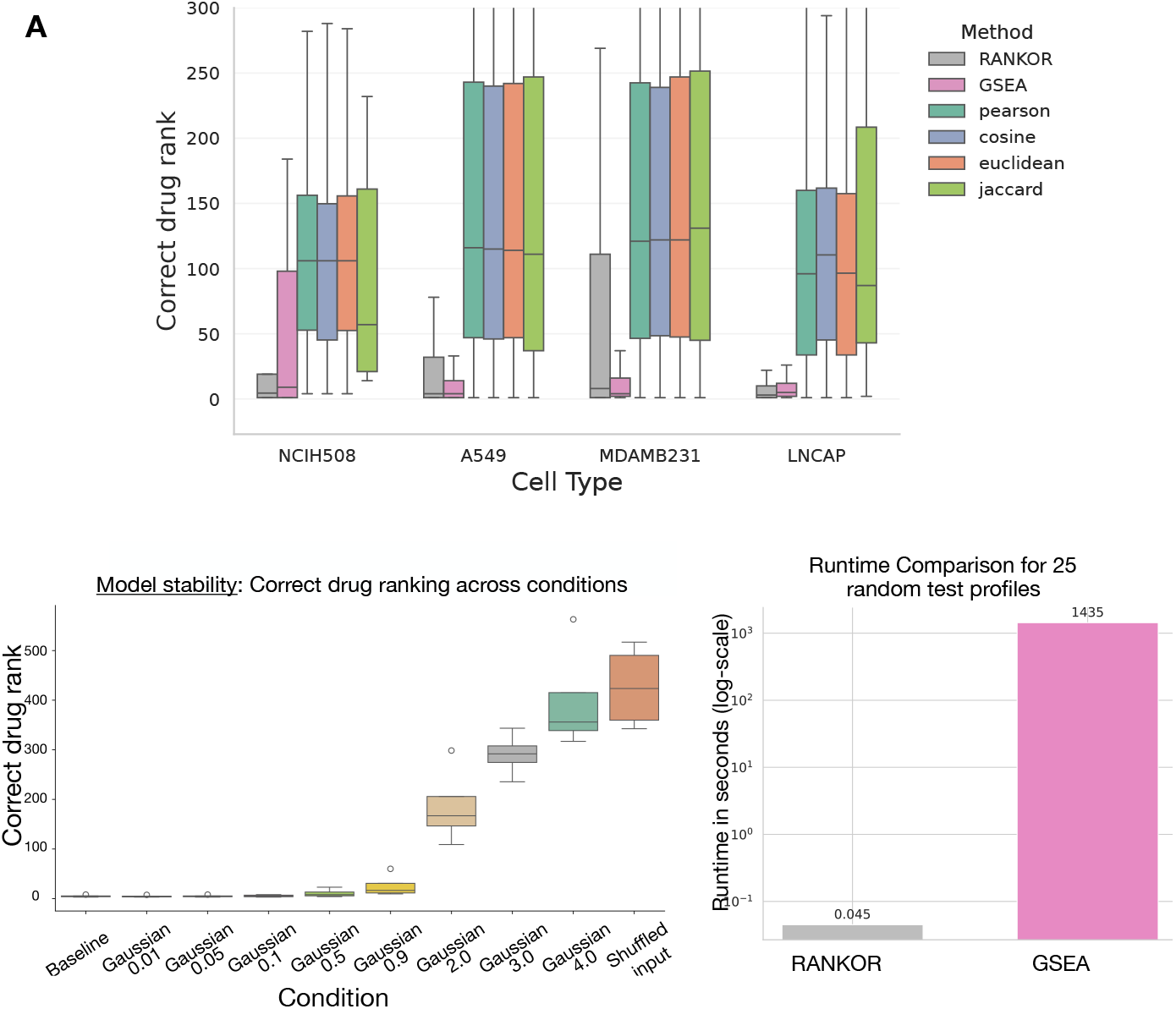
Robustness, negative control, and computational efficiency of RANKOR. (A) Drug ranking performance across four cell lines (NCIH508, A549, MDAMB231, LNCAP) comparing RANKOR with GSEA and similarity-based baselines (Pearson, cosine, Euclidean, Jaccard) (B) Performance under increasing noise and shuffled input.(C) Runtime comparison across randomly sampled signatures (seconds in log10 scale).

Across all cell types, RANKOR consistently achieved lower rankings compared to classical similarity methods and competitive median ranks compared to alternative statistical approaches, such as GSEA. These results demonstrate improved robustness and generalization when drug ranking is learned directly in a structured latent space rather than inferred from direct signature matching.

### 2.5 Robustness to transcriptomic perturbations and negative control analysis

To evaluate robustness to input perturbations, we applied progressively increasing Gaussian noise to gene expression profiles prior to drug ranking (Fig. 3B). Under mild perturbations (noise *≤*0.1), RANKOR maintained stable drug rankings, indicating resilience to realistic fluctuations in gene-level signal. With increasing noise magnitude, RANKOR exhibited a gradual and monotonic degradation in performance. RANKOR operates in a learned continuous latent space that captures fine-grained transcriptomic structure, rendering it more sensitive to substantial perturbations while enabling higher discriminative precision when biologically meaningful signal is preserved.

To further validate that performance depends on structured transcriptomic information rather than statistical artifacts, gene expression values were randomly permuted within each signature. This shuffling destroys gene-order structure while preserving the underlying distributions. These results confirm that predictive performance arises from structured biological signal. All together, the perturbation and shuffle analyses demonstrate that RANKOR achieves robust performance under realistic noise while retaining higher-resolution sensitivity to transcriptomic structure beyond pathway-level aggregation.

### 2.6 Runtime performance and scalability analysis

The machine learning nature of RANKOR enables computationally efficient predictions. We benchmarked per-signature inference time for RANKOR and GSEA across 25 randomly sampled test profiles. Runtime was measured end-to-end for each method, excluding one-time model and library loading (approx. 0.01 second). For RANKOR, this included latent embedding inference and cosine similarity–based drug ranking across the candidate compound space. For GSEA, timing encompassed pre-ranked enrichment analysis against the LINCS consensus perturbation signatures followed by concordance-based drug ranking. Across test signatures, RANKOR demonstrated substantially lower inference times (Fig. 3C), achieving orders-of-magnitude faster per-query execution compared to GSEA. This efficiency advantage reflects the lightweight forward-pass architecture of RANKOR relative to permutation-based enrichment procedures, enabling rapid large-scale transcriptomic drug prioritization without sacrificing predictive performance.

### 2.7 Drug prioritization for transcriptionally unseen compounds

A key limitation of traditional signature reversal approaches such as enrichment-based and connectivity-mapping methods, including GSEA or correlation-based connectivity scoring, is that they require reference transcriptional profiles for each compound (Subramanian et al. 2005, 2017; Duan et al. 2016). As a result, drugs lacking perturbational signatures cannot be evaluated within these frameworks. In contrast, we built RANKOR to leverage the shared pharmacological structure of the chemical latent space to enable prioritization beyond the experimentally profiled drug universe.

To evaluate whether therapeutics that have never been transcriptionally profiled can be correctly prioritised, we designed an unseen-drug evaluation setting in which entire compounds were excluded from transcriptomic training. In this scenario, no gene expression signatures associated with these drugs were available to the model. Instead, unseen compounds were incorporated into the prioritization framework exclusively through their chemical structure.

Specifically, canonical SMILES representations of test drugs, which are held out from the NN1 and NN2 training, were transformed into molecular fingerprints and projected into the chemical latent space using the pretrained chemical encoder. This procedure yields a latent embedding for each unseen drug, enabling its inclusion in the ranked candidate list despite the absence of perturbational transcriptomic data. Consequently, RANKOR can prioritize compounds that have never been experimentally screened in transcriptional assays or disease-relevant cellular contexts.

Figure 4A summarizes prioritization performance across unseen compounds. We report examples of best-and worst-performing drugs across all cross-validation folds and cellular contexts. As shown in Fig. 4A, RANKOR maintains a good recovery of the correct compound for a substantial fraction of unseen drugs, indicating that cross-space alignment successfully transfers mechanism-level information from transcriptomic to chemical representations.

**Fig. 4.**
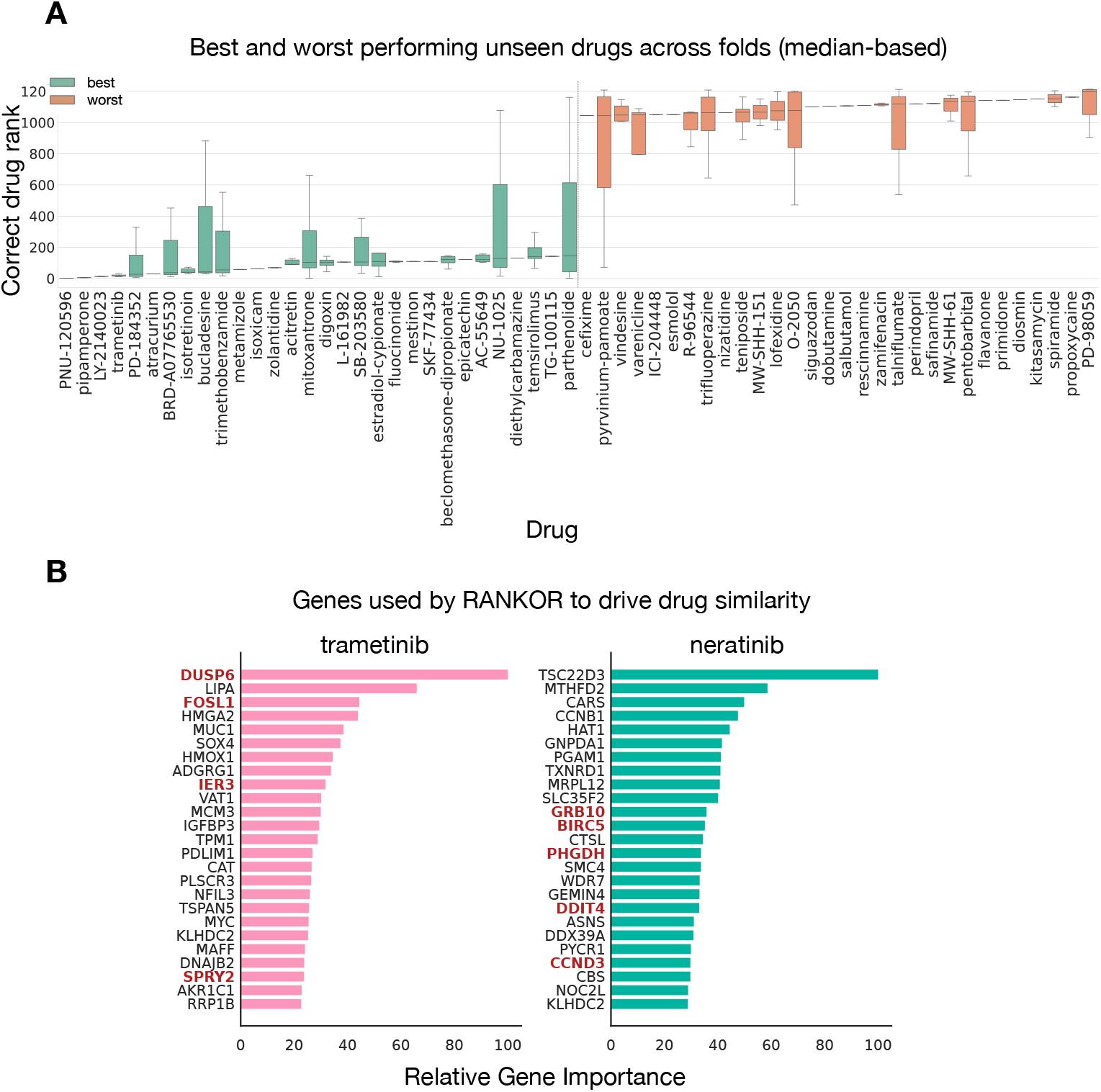
Performance on transcriptionally unseen drugs and biological interpretability of RANKOR. **(A)** Best- and worst-performing unseen drugs across cross-validation folds (median rank shown). RANKOR successfully prioritizes a substantial subset of compounds despite the absence of prior transcriptional profiling. **(B)** Gene-level attribution analysis for trametinib and neratinib as it is calculated by mean over all samples of all cell lines. The most influential genes driving similarity reflect known mechanism-of-action biology. For trametinib, MAPK pathway–associated genes (e.g., DUSP6, SPRY2, FOSL1) are highlighted in red. For neratinib, ERBB signaling and downstream proliferation regulators (e.g., GRB10, BIRC5, PARP1, CCND3, PHGDH, DDIT4) dominate. Relative gene importance was computed via gradient-based attribution from transcriptomic input to drug similarity score.

Performance variability across compounds reflects biological and pharmacological factors rather than model instability. Drugs inducing strong and distinctive transcriptional programs are more readily aligned to their chemical embeddings, whereas compounds producing weak, context-dependent, or highly varying responses are inherently more difficult to prioritize. Notably, when the exact compound is not ranked first, the model frequently prioritizes pharmacologically related agents with shared pathway activity or target class, supporting the biological coherence of the learned chemical space. Representative examples of prioritization outcomes for transcriptionally unseen compounds are shown in Table 1.

**Table 1.** Examples of prioritization outcomes for transcriptionally unseen drugs. When the correct compound is not ranked first, RANKOR frequently prioritizes mechanistically related agents, reflecting shared pathway activity captured in the chemical latent space.

| Correct drug | Most frequently confused with | Mechanistic relationship |
| --- | --- | --- |
| Trametinib | U-0126 (MEK1/2 inhibitor) | Both compounds inhibit the MAPK/ERK signaling cascade through MEK blockade, producing highly concordant transcriptional responses (Yamaguchi et al. 2011; Little et al. 2011). |
| Pipamperone | Haloperidol (D2 receptor antagonist) | Shared dopamine D2 receptor antagonism within the antipsychotic drug class; strong pharmacological and transcriptional similarity is expected (Seeman 2002; Lieberman et al. 2005). |
| PNU-120596 | Articaine | Limited structural similarity; prioritization likely reflects partial overlap in neuronal excitability and synaptic signaling programs rather than direct target congruence (Hurst et al. 2005; Kress et al. 2008). |

### 2.8 Drug prioritisation relies on biologically meaningful transcriptional programs

Beyond ranking performance, we examined whether RANKOR bases its decisions on mechanistically relevant genes rather than spurious transcriptomic patterns. To this end, we performed gradient-based attribution analysis to quantify the contribution of each gene to the similarity score between transcriptomic signatures and candidate drug embeddings.

For a given compound, gradients were computed with respect to the cosine similarity between the projected transcriptomic embedding and the target chemical embedding. Gene-level importance was estimated by averaging absolute gradient contributions across signatures, scaled by input expression values (saliency weighting). This procedure yields a ranked list of genes most influential in driving drug similarity.

As shown in Fig. 4B, attribution profiles align closely with known mechanisms of action. For trametinib, a selective MEK1/2 inhibitor targeting the MAPK/ERK signaling cascade (Kidger et al. 2018; Hatzivassiliou et al. 2012), highly ranked genes include *DUSP6, SPRY2, FOSL1*, and other MAPK-responsive transcriptional regulators. These genes are canonical downstream effectors and feedback modulators of ERK signaling, consistent with trametinib-induced pathway inhibition.

For neratinib, an irreversible pan-ERBB tyrosine kinase inhibitor targeting HER2/ERBB2 and EGFR that suppresses downstream PI3K–AKT and MAPK signaling pathways (Rabindran et al. 2004; Collins et al. 2019), the most influential genes include *GRB10, BIRC5, PARP1*, and the cell-cycle regulator *CCND3*, together with metabolic and stress-response genes such as *PHGDH* and *DDIT4*. These genes have previously been associated with ERBB signaling, apoptosis regulation, cell-cycle progression, and metabolic adaptations in HER2-driven tumors (Holt and Siddle 2005; Ryan et al. 2006; Nowsheen et al. 2012; Locasale et al. 2011). Their identification among the most influential features suggests that the model captures biologically meaningful transcriptional programs associated with ERBB pathway inhibition.

Importantly, the attribution results indicate that RANKOR prioritizes drugs based on coherent pathway-level transcriptional programs consistent with established pharmacology. The model does not rely on arbitrary gene patterns but instead leverages biologically grounded signals aligned with known drug targets and downstream regulatory networks.

### 2.9 Prediction of applied drug in real patient T-ALL single-cell data

While the previous analyses were performed on cell line perturbation datasets—reflecting the environment in which RANKOR was trained, the key translational question is whether the framework generalizes to patient-derived data. To address this, we evaluated the ability of RANKOR to recover therapeutically relevant compounds in a T cell acute lymphoblastic leukemia (T-ALL) single-cell dataset comprising malignant and healthy T cell populations (Caron et al. 2020). The different cell types found in the patient data are depicted in Fig. 5A. After preprocessing, T cells were isolated and subclustered using Leiden community detection to capture transcriptionally distinct cellular states within the malignant compartment (Fig. 5B).

These subclusters were subsequently annotated into major T cell subtypes, including naive/memory-like T cells, cytotoxic T cells, and immature precursor T cells, based on cell type specific marker genes. The naive/memory-like cluster was characterized by the expression of genes associated with quiescent and homeostatic T cell states, including IL7R, TXNIP, NFKBIA, PIK3IP1, and IFITM1, consistent with central memory or resting T cells. The cytotoxic cluster exhibited strong enrichment of effector molecules and cytolytic machinery, including NKG7, GZMA, GZMK, CCL5, CST7, and SRGN, along with differentiation markers such as ID2, KLRG1, and KLRB1, indicative of activated cytotoxic or NK-like T cells. In contrast, the immature precursor cluster was defined by genes associated with early T cell development and proliferative programs, including SOX4, DNTT, MYB, BCL11A, CDK6, CD99, TCF7, and STMN1, consistent with thymocyte-like or leukemic precursor states.

**Fig. 5.**
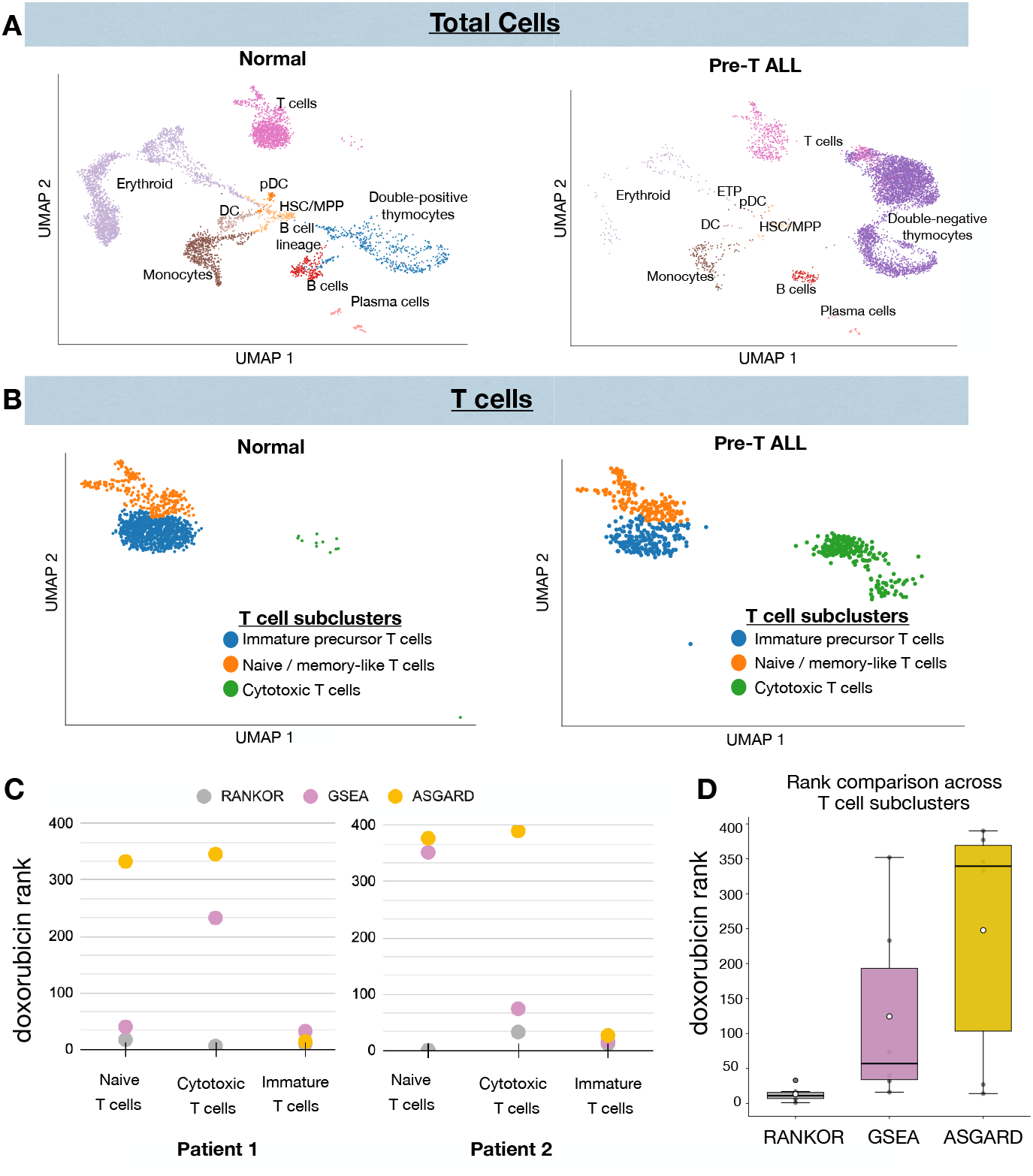
Drug prioritization in T-ALL single-cell clusters. (A) UMAP visualization of T cell populations from healthy donors and T-ALL patients, showing clustering of healthy T cells and malignant T cell populations. Patients have been treated with high dosage doxorubicin. (B) UMAP of the T cell subclusters. The T-cells were isolated and the subclusters were annotated in order to apply different methods on each of the subtypes individually and to leverage the full potentials of single cell data. (C) Cmparison of doxorubicin ranking across clusters for RANKOR, GSEA, and ASGARD for the two patients. Lower rank indicates better prioritization (y-axis). (D)Comparison of rankings across T cell subclusters.

For each patient-specific cluster, standardized transcriptional signatures were computed relative to matched healthy T cell populations and shortened according to the LINCS landmark gene space, enabling direct comparison with perturbational reference profiles. All patients were treated according to the DFCI protocol, which includes high-dose doxorubicin.

Across subtypes and patients, doxorubicin was consistently ranked among the top candidates by RANKOR, indicating strong predicted power of leukemia-associated transcriptional programs across diverse T cell states (Fig. 5C and Fig. 5D). In contrast, baseline approaches based on the GSEA and the single-cell drug repurposing framework ASGARD (He et al. 2023) produced weaker prioritization, assigning higher (less favorable) ranks to doxorubicin across multiple subtypes. This pattern reflects a strong dependency on cellular state, with the immature precursor cluster showing consistently high detection across all methods due to its proliferative signature, in line with doxorubicin’s mechanism of action. In contrast, naive-like and cytotoxic clusters exhibit more complex transcriptional programs, where enrichment-based approaches often fail, assigning unfavorable ranks. RANKOR, however, maintains robust prioritization across all subtypes and patients, indicating its ability to capture structured transcriptional signals beyond simple gene set overlap.

Overall, these results suggest that RANKOR effectively identifies clinically relevant therapeutic signatures across 6 heterogeneous T cell subpopulations from 2 patients.

#### Subtype-specific transcriptional programs driving doxorubicin similarity

To interpret subtype-specific drug predictions, we applied gradient-based attribution to identify the genes most strongly contributing to doxorubicin similarity within each T cell subtype (Fig. 6). Naive / memory-like T cells show contributions from genes such as ARPP21, FXYD2, and SMPD3, indicating subtype-specific regulatory and metabolic programs. Cytotoxic T cells are driven by signaling and regulatory genes including ZNF816, TNFRSF21, DAG1, and CDKN2A, consistent with an activated or stress-responsive state, in line with the induction of DNA damage and cell-cycle arrest programs by anthracyclines such as doxorubicin (Carvalho et al. 2009; Thorn et al. 2011). In contrast, immature precursor T cells exhibit contributions from genes such as B2M, CD52, DDIT4, and HERPUD1, reflecting transcriptional programs associated with developmental and stress-related processes, including mTOR and endoplasmic reticulum stress responses that are known to be activated following chemotherapy-induced cellular stress (Yang et al. 2014).Importantly, the above mentioned genes differ from those used for subtype annotation, reflecting transcriptional features relevant to drug response rather than cell identity These findings are further consistent with the presence of immature thymocyte-like populations in T cell acute lymphoblastic leukemia, which exhibit distinct transcriptional programs linked to proliferation and stress adaptation (Belver and Ferrando 2016; Coustan-Smith et al. 2009).

**Fig. 6.**
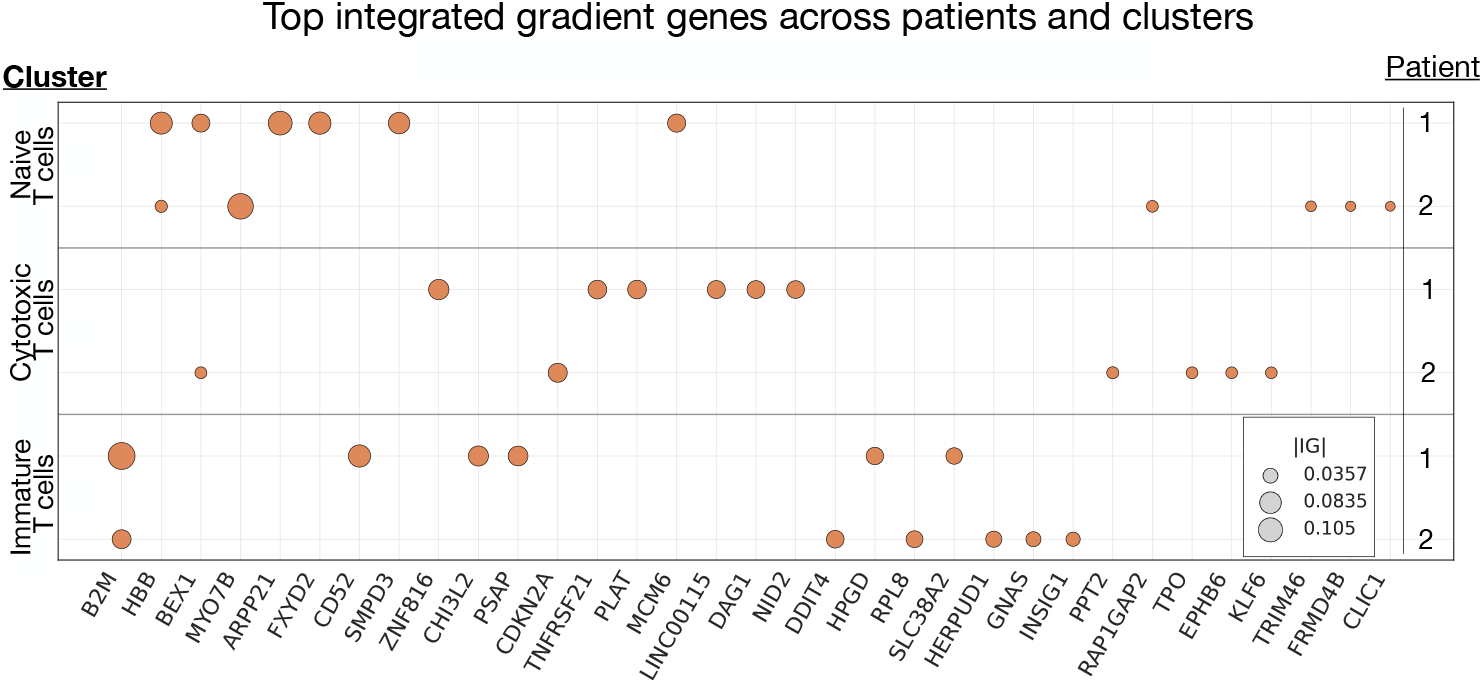
Top selected genes for RANKOR prioritization of doxorubicin in different single cell clusters across patients. The selection is based on integrated gradient-based attribution. Dot size indicates importance (absolute attribution).

### 2.10 Glioblastoma single-cell analysis reveals subtype-resolved drug prioritization

As an additional single cell application with real patient data, we applied RANKOR to recover clinically relevant compounds in a highly heterogeneous solid tumor setting. Specifically, we analyzed single-cell RNA-seq profiles from patient-derived glioblastoma slice cultures with matched vehicle- and etoposide-treated samples across multiple patients (Cheng et al. 2024). Following preprocessing and batch integration, we identified non-malignant microenvironmental populations using established lineage markers, and classified malignant cells as transformed tumor cells based on the characteristic glioblastoma-associated imbalance of chromosome 7 gain and chromosome 10 loss, in line with previously described single-cell GBM annotation strategies (Neftel et al. 2019). We then projected transformed cells into glioblastoma cellular programs defined by the framework of Neftel et al., assigning cells to mesenchymal-like (MES-like), astrocyte-like (AC-like), oligodendrocyte progenitor-like (OPC-like), and neural progenitor-like (NPC-like) programs, while also capturing proliferative G1/S and G2/M programs as separate malignant states. This yielded a subtype-resolved view of intra-tumoral heterogeneity across patients.

Analysis of variance of single cell expression (Fig. 7A) illustrates that the dataset contained both inter-patient variation and substantial within-patient cellular diversity, with transformed and non-transformed populations separable and the malignant compartment further decomposable into distinct glioblastoma programs. Control cells and cells after application of the drug etoposide (Fig. 7B) are distributed across the different states of all cell types(Fig. 7C). Notably, vehicle- and etoposide-treated cells remain broadly distributed across the same similar regions in the displayed latent space rather than separating into a single treatment-specific cluster, supporting the idea that drug response must be evaluated in a state-specific rather than globally averaged manner. This is consistent with the known transcriptional plasticity of glioblastoma, where malignant cells occupy recurrent cellular states whose relative abundance varies across tumors (Wang et al. 2022).

**Fig. 7.**
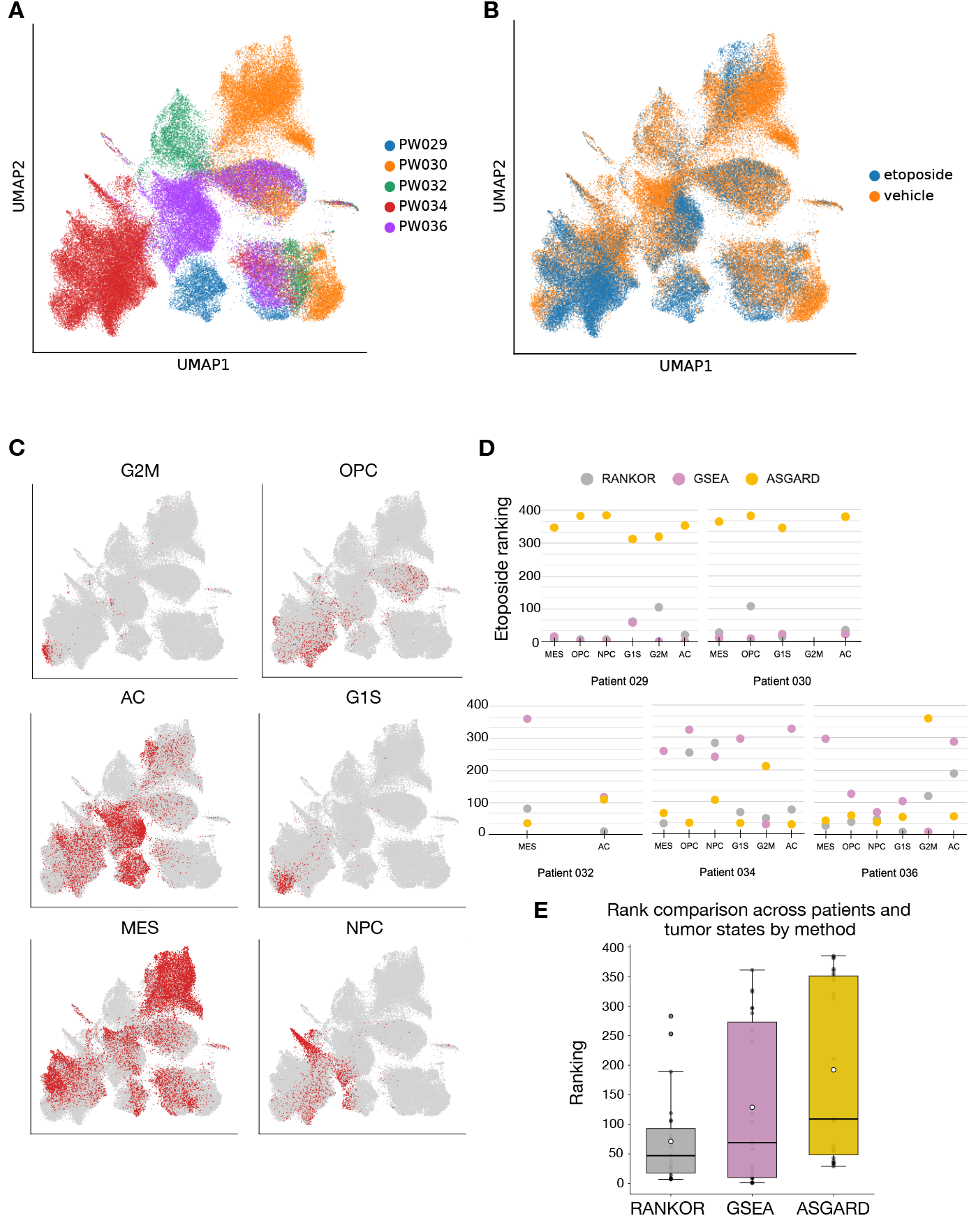
Subtype-resolved drug prioritization in glioblastoma single-cell data. (A) UMAP visualization of individual cells, where each point represents a single cell and colors indicate the patient of origin. Cells positioned closer together exhibit more similar transcriptional profiles, allowing visualization of patient-specific and shared cellular populations. (B) Cells colored by treatment condition (vehicle vs. etoposide). (C) Malignant glioblastoma cells were further grouped into transcriptional cell states and cells belonging to a group colored in red, including MES (mesenchymal-like), AC (astrocyte-like), OPC (oligodendrocyte progenitor-like), NPC (neural progenitor-like), and proliferative states associated with the cell cycle (G1/S and G2/M). (D) Etoposide ranking across patients and cell states, comparing RANKOR, GSEA, and ASGARD. Lower ranks indicate better recovery of the true perturbagen. (E) Comparison of rankings across patients and tumor states for all methods.

We next constructed patient- and subtype-specific pseudo-bulk signatures by contrasting each treated population against the matched vehicle baseline within the same patient and cell state, and used these signatures for drug prioritization. Etoposide is a topoisomerase II inhibitor that induces DNA damage and is most active in proliferative cells. The recovery of etoposide provides a biologically meaningful test of whether the models capture treatment-relevant transcriptional structure.

In Fig. 7D, we compare etoposide rankings across malignant glioblastoma cell states as they are. Notably, mean ranking performance differed across methods, with RANKOR achieving the lowest overall mean rank, followed by GSEA, while ASGARD showed substantially higher ranks (Fig. 7E). These results indicate that RANKOR provides more consistent prioritization performance across glioblastoma cell states, while GSEA remains comparatively competitive but exhibits reduced overall rankings.

Importantly, performance varied across patients and cellular states, underscoring the biological complexity of glioblastoma drug response. In some malignant subtypes, all methods recovered etoposide well, whereas in others the rankings varied substantially across patients. ASGARD generally showed weaker recovery, while GSEA exhibited higher variability across patient-specific cellular contexts. Differences in etoposide ranking across malignant cell states suggest that the transcriptional response to the drug is not equally preserved across all tumor subpopulations, potentially reflecting heterogeneous drug sensitivity among glioblastoma cell states. In this setting, RANKOR identified etoposide across several malignant subpopulations, in some cases with lower (better) rankings and in others with performance comparable to the best-performing method, demonstrating the utility of subtype-resolved single-cell analyses for investigating drug responses across distinct tumor cell populations.

## 3 Methods

### 3.1 Bulk data processing

Transcriptomic perturbation profiles were obtained from the LINCS L1000 resource (Subramanian et al. 2017). The L1000 assay directly measures 978 landmark genes and infers the expression of additional genes using models trained on large-scale transcriptomic data. Consistent with prior work (Katsaouni and Schulz 2026), we used the full set of measured and inferred genes to capture comprehensive transcriptional responses (Jiang et al. 2024).

We used Level-5 consensus signatures, where each signature represents a specific compound–cell line–dose–time condition and is defined as the median z-score differential expression across biological replicates. Signatures were downloaded from the CLUE platform (https://clue.io). Each signature is associated with a Transcriptional Activity Score (TAS), which summarizes both the magnitude of differential expression and replicate consistency. Following established practice, we retained signatures with TAS *≥*0.2 and excluded compounds represented by fewer than 10 signatures (Huang et al. 2025).

Mechanism-of-action (MoA) annotations were obtained from curated sources (Huang et al. 2018) and aligned with recent transcriptomics-based MoA prediction studies (Jiang et al. 2024). After filtering, the dataset comprised 28,769 transcriptional signatures from 1,214 compounds and was used to learn the transcriptomic latent space.

### 3.2 Generalization aware dataset splitting

To ensure realistic assessment of generalization and to avoid information leakage, data were split at either the compound or cell-type level, following the strategy described in our previous manuscript on transcriptomic representation learning (Katsaouni and Schulz 2026). In compound-based splits, complete compounds were withheld from training to evaluate performance on unseen drugs within known mechanisms of action. Specifically, 10% of the compounds were kept out of the training for each of the folds. In cell-type–based splits, all signatures from selected cell lines were excluded during training to assess transfer to new cellular contexts. The held-out lines were NCIH508 (colorectal adenocarcinoma; 26 signatures), A549 (lung carcinoma; 2,209 signatures), MDAMB231 (triple-negative breast cancer; 646 signatures), and LNCaP (prostate adenocarcinoma; 94 signatures), representing diverse tissue origins (Katsaouni and1 Schulz 2026). Together, these complementary splitting schemes enforce strict separation between training and evaluation data across both chemical and biological dimensions.

### 3.3 Chemical structure representation and preprocessing

Chemical structures corresponding to LINCS compounds were obtained in SMILES format (O’Boyle 2012). To derive fixed-length chemical representations, each SMILES string was converted into a binary Morgan fingerprint using a circular fingerprinting scheme. Fingerprints were generated with a radius of 2 and a dimensionality of 256 bits, providing a compact representation of local chemical substructures.

These chemical fingerprints served as input to the chemical encoder and were used to construct the chemical latent space during RANKOR learning. Each compound is represented by a single chemical embedding, enabling alignment between transcriptional responses and chemical structure in downstream drug prioritization.

### 3.4 Hierarchical transcriptomic latent space

To capture biologically meaningful structure in drug-induced transcriptional responses, we leveraged a hierarchical transcriptomic latent space learned using contrastive representation learning. The construction and validation of this latent space are described in detail in our companion manuscript (Katsaouni and Schulz 2026); here we provide a concise summary relevant to the RANKOR framework.

Briefly, transcriptomic signatures were embedded using a neural encoder (denoted as NN1 in Fig. 1) trained to jointly organize profiles at two biological levels: mechanism of action (MoA) and individual compound. The NN1 gets as input the z-scores of the transcriptomic signatures and outputs a 256-dimensional latent vector *z*_x_ ∈ℝ^256^, where x referes to each of the signatures. The model employs a shared encoder followed by two cooperative classification objectives, encouraging signatures from drugs with the same MoA to form coherent clusters, while preserving compound-specific substructure across doses, time points, and cellular contexts. This hierarchical organization reflects known pharmacological relationships, with compact MoA-level groupings and finer-grained drug-level separation.

Angular margin–based supervision was used to enforce separation between MoAs in the latent space, while a secondary objective grouped replicate signatures of the same compound. Together, these objectives yield a structured embedding in which transcriptional responses are organized according to shared biological mechanisms while retaining meaningful within-mechanism variability. The resulting latent space serves as the transcriptomic foundation for linking gene expression signatures to chemical representations in downstream drug prioritization.

Model training and quantitative evaluation of MoA separation and compound-level organization are described in detail in (Katsaouni and Schulz 2026).

### 3.5 Chemical reference latent space

In parallel to the transcriptomic representation, we constructed a chemical latent space in which each compound is represented by a single fixed-length embedding derived solely from its molecular structure. This chemical latent space serves two critical purposes within RANKOR. First, it provides a unique, one-point-per-drug representation that enables the alignment of multiple transcriptomic signatures associated with the same compound into a unified chemical coordinate. Second, it enables drug prioritization for previously unseen compounds that lack transcriptional profiling, by mapping chemical structure directly into the latent space.

#### Chemical input representation

For each compound, canonical SMILES strings were obtained from LINCS metadata. SMILES entries marked as restricted or failing RDKit parsing were excluded. Each valid SMILES string was converted into a binary Morgan fingerprint using RDKit (`GetMorganFingerprintAsBitVect`) with radius *r* = 1 and dimensionality of 256 bits. To ensure a unique chemical representation per compound, a single fingerprint was retained for each unique drug, yielding one binary feature vector per drug.

#### Chemical encoder architecture

Chemical fingerprints were embedded using a feed-forward neural network that maps the 256-bit binary input into a 256-dimensional continuous latent embedding. The encoder consists of two hidden layers of size 512 with ReLU activations, followed by a linear output layer:

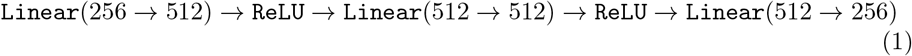

The output of the chemical encoder is a 256-dimensional latent representation

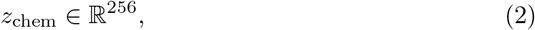

which corresponds to the learned chemical embedding of the input compound. The resulting embeddings were subsequently *𝓁*_2_-normalized prior to loss computation.

#### Mechanism-aware supervision with ArcFace

To structure the chemical latent space according to pharmacological mechanism, the chemical encoder was trained using an ArcFace classification objective (Deng et al. 2019) over mechanism-of-action (MoA) labels. ArcFace enforces angular separation between classes in normalized embedding space by introducing an additive angular margin to the target class.

Given a normalized chemical embedding **z**_*i*_ and normalized class weight vector 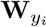 for compound *i* with MoA label _*yi*_, the cosine similarity is defined as:

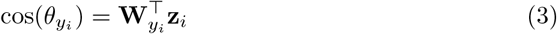

An additive angular margin *m* = 0.2 was applied to the target class, and logits were scaled by a factor *s* = 30. This objective encourages compounds sharing the same MoA to form compact clusters in angular space while remaining well separated from compounds acting through different mechanisms.

The chemical encoder and ArcFace head were trained jointly using the Adam optimizer with learning rate 10^*−*4^ and batch size 128. Training was performed for multiple epochs until convergence. Only chemical fingerprints and MoA labels were used during training; no transcriptomic information was provided to the chemical encoder. The resulting model produces a single, mechanism-aware embedding for each compound based solely on chemical structure.

### 3.6 Cross-space modeling: mapping transcriptomic to chemical space

To connect transcriptional responses to chemical structure and enable direct drug prioritization, we learned a cross-space mapping model (NN3) that projects points from the transcriptomic latent space into the chemical latent space. This model is central to RANKOR for two reasons. First, it aligns multiple transcriptomic signatures associated with the same drug (across dose, time, and cell type) toward a single chemical embedding, thereby linking experimental variability in transcriptional space to a unique compound representation. Second, because chemical embeddings can be computed for any drug from its structure, the mapping enables prioritization against a candidate set that includes compounds without transcriptional profiling.

#### Training pairs and targets

For each fold, we constructed paired training examples by matching transcriptomic signatures to their corresponding compound identity. Transcriptomic inputs were obtained by embedding each signature with the transcriptomic encoder (NN1), yielding a 256-dimensional vector **z**^x^ ∈ℝ^256^. The corresponding targets were 256-dimensional chemical embeddings **z**^chem^ ∈ℝ^256^ computed from Morgan fingerprints using the chemical encoder (NN2). Training was restricted to drugs with valid SMILES and available embeddings in both spaces. Importantly, multiple transcriptional signatures can map to the same target drug embedding, reflecting different experimental conditions for the same compound.

#### Mapping network architecture

The cross-space model is a fully connected regression network that maps transcriptomic embeddings to chemical embeddings:

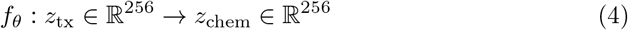

We implemented *f*_*θ*_ as a multilayer perceptron with two hidden layers (512 units each) and ReLU activations:

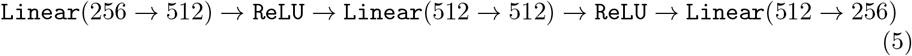

The output 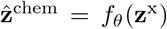 represents the predicted location in chemical latent space corresponding to the observed transcriptional signature.

We trained the mapping model to maximize angular agreement between predicted and target chemical embeddings. Specifically, we minimized a cosine-distance loss:

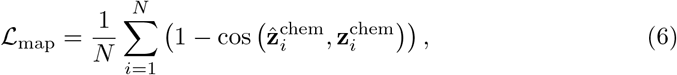

where cos(*·,·*) denotes cosine similarity. Model parameters were optimized, using a grid search, with Adam (learning rate 10^*−*3^) for 50 epochs using mini-batches of size 64. A separate mapping model was trained for each fold to match the corresponding transcriptomic and chemical encoders.

All models were implemented in PyTorch and trained on GPU. Chemical embeddings were computed using fold-specific ArcFace chemical encoders, and transcript embeddings were computed using fold-specific transcriptomic encoders.

#### Drug ranking by similarity in chemical space

At inference time, a query signature (e.g., disease vs. healthy, or pre-vs. post-treatment) is first embedded into the transcriptomic latent space using NN1 and then projected into chemical space using NN3 to obtain 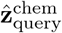. Candidate drugs are ranked by cosine similarity between 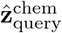 and each drug embedding in the chemical latent space. The resulting output is an ordered list of candidate compounds, enabling direct prioritization without reconstructing high-dimensional gene expression profiles.

### 3.7 Stability and robustness to input perturbations

To assess the robustness of RANKOR to perturbations in transcriptomic data, we performed a systematic stability analysis across all cross-validation folds. For each fold, model performance was evaluated under increasing levels of additive Gaussian noise as well as under gene-shuffling perturbation applied to the test-set gene expression matrix.

#### Noise injection

Let **X**_test_ ∈ℝ^*n×g*^ denote the test gene expression matrix, where *n* is the number of signatures and *g* the number of genes. For each noise level *σ*, Gaussian noise was added independently to each gene expression value:

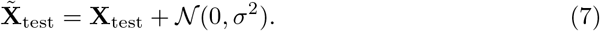

Noise scales *σ* ∈ {0.01, 0.05, 0.1, 0.5, 0.8, 0.9, 2.0, 3.0, 4.0} were evaluated to simulate increasing degrees of measurement variability and signal degradation.

For each noise level and fold the noisy gene expression matrices were embedded into the transcriptomic latent space using the trained transcript encoder (NN1). The noisy embeddings were projected into chemical latent space using the trained cross-space mapping network (NN3). Afterwards, drug prioritization was performed by computing cosine similarity between predicted chemical embeddings and all reference drug embeddings and for each test signature, we recorded the rank of the correct compound among all candidates.

All results were recorded separately per fold and aggregated across folds to evaluate consistency. This analysis quantifies how sensitive RANKOR’s prioritization performance is to perturbations in input gene expression values and provides a measure of model stability under increasing noise.

#### Gene shuffling analysis

To further evaluate whether RANKOR relies on gene-specific biological signal rather than global distributional properties, we performed a gene shuffling control experiment. For each test signature, gene expression values were randomly permuted across gene features while preserving the overall distribution of expression values within each sample.

Formally, let **x**_*i*_ ∈ ℝ^*g*^ denote the expression vector for signature *i*. A shuffled profile 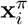 was generated by applying a random permutation *π* over gene indices:

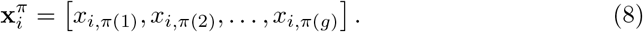

This procedure maintains the distribution of expression values but disrupts gene-level correspondence and consequently pathway structure.

Shuffled signatures were then processed through the full RANKOR pipeline, including transcriptomic embedding, cross-space projection, and drug ranking.

This analysis isolates the contribution of gene-specific transcriptional patterns to prioritization performance and serves as a stringent negative control for biological signal utilization.

### 3.8 Model explainability and gene attribution

To investigate whether RANKOR prioritization is driven by biologically meaningful transcriptional programs, we performed a gradient-based explainability analysis to identify genes contributing to drug-specific similarity predictions.

We constructed a model linking gene expression input directly to chemical latent representations by sequentially combining the transcriptomic encoder (NN1) and the cross-space mapping network (NN3). This end-to-end differentiable architecture enabled attribution of drug similarity scores to individual gene features.

For a given drug *d*, we quantified gene importance by computing gradients of the cosine similarity between predicted drug embeddings and the reference chemical embedding of *d*. Formally, let **x**_*i*_ ∈ℝ^*g*^ denote the transcriptomic profile of sample *i*, and let 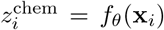 represent the predicted drug embedding produced by the full RANKOR model. Let 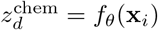 denote the reference chemical embedding of drug *d*. Drug similarity is defined as:

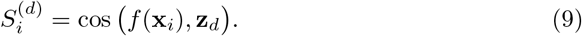

Gene importance scores were computed as the gradient of this similarity with respect to each gene:

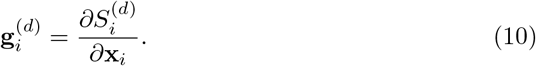

To improve interpretability, we employed a saliency-weighted formulation in which gradients were multiplied by the input expression values:

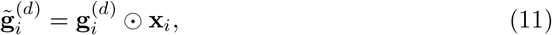

where ⊙ denotes element wise multiplication. This formulation highlights genes whose expression magnitude and sensitivity jointly influence similarity predictions.

Gene attribution scores were averaged across all transcriptional signatures associated with drug *d*:

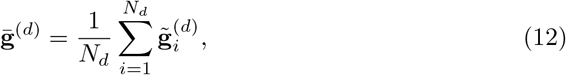

where *N*_*d*_ is the number of signatures linked to drug *d*.

Genes were ranked by the absolute magnitude of their mean attribution scores, and the top contributing genes were retained for downstream interpretation and visualization.

### 3.9 Single-cell transcriptomic datasets and preprocessing

To evaluate the applicability of RANKOR to patient-derived single-cell data, we analyzed two publicly available single-cell RNA sequencing cohorts representing distinct disease and treatment contexts.

#### Acute leukemia cohort

Single-cell transcriptomic profiles were obtained from the pre–T acute lymphoblastic leukemia dataset GSE132509 by (Caron et al. 2020), comprising bone marrow mononuclear cells from leukemia patients treated under the Dana–Farber Cancer Institute (DFCI) protocol, characterized, among others, by high cumulative doxorubicin exposure (300 mg/m^2^) (Moghrabi et al. 2007). We analyzed samples from two leukemia patients alongside three healthy peripheral blood mononuclear cell controls. Raw count matrices were loaded from matrix market files and combined into a unified AnnData object with harmonized gene annotations and cell metadata.

#### Glioblastoma cohort

We further analyzed the glioblastoma single-cell dataset GSE148842 (Cheng et al. 2024), focusing on five patients profiled under ex vivo drug perturbation. For each patient, matched vehicle-treated and etoposide-treated samples were included. Count matrices were imported from compressed transcript count tables, converted to sparse matrices, and merged across patients while preserving sample and treatment annotations.

#### Single-cell data processing

For both datasets, preprocessing was performed using Scanpy (Wolf et al. 2018). Cells expressing fewer than 200 genes or exhibiting high mitochondrial transcript proportions (*>* 10%) were excluded. Gene features expressed in very few cells were removed. Count matrices were library-size normalized to 10,000 counts per cell and log-transformed.

Where multiple samples or patients were combined, batch effects were corrected using Harmony (Korsunsky et al. 2019) integration on principal component embeddings. Neighborhood graphs and UMAP projections were computed from corrected representations. Cell-type annotation was performed using CellTypist (Xu et al. 2023), enabling harmonized immune cell labeling across disease and control samples.

To derive drug/disease-relevant transcriptional signatures, cells were stratified by condition (e.g., leukemia vs. healthy; drug-treated vs. vehicle) and, where appropriate, further subset by annotated cell type. T cell populations were isolated and subclustered using Leiden community detection to capture disease-specific cellular states.

Transcriptional signatures were derived by computing standardized z-score profiles between matched conditions, generating perturbation signatures directly compatible with the RANKOR input framework.

To ensure compatibility with the perturbational reference space, single-cell expression matrices were aligned to the LINCS landmark gene set. Genes absent from the single-cell datasets were introduced as zero-expression features, preserving gene ordering and dimensional consistency with the transcriptomic encoder. The resulting signatures were used as query inputs for drug prioritization.

### Baseline: Gene set enrichment–based drug prioritization

To benchmark RANKOR against enrichment-based approaches, we used the Gene Set Enrichment Analysis (GSEA) pipeline using the LINCS L1000 perturbational reference collection. For each test sample, standardized gene expression signatures (z-scores) were used to construct a ranked gene list, which served as input to a GSEA analysis implemented via the GSEApy Python package (Fang et al. 2023).

As a reference database, we used the *LINCS L1000 Chemical Perturbation Consensus Signatures* collection, which contains consensus transcriptional profiles for small-molecule perturbations across multiple cell lines. Enrichment scores were computed separately for upregulated and downregulated perturbational signatures using 100 permutations.

For each perturbagen, enrichment results were parsed to extract drug identity and directionality (up/down signatures). The most significant enrichment result per drug was retained separately for upregulated and downregulated signatures based on p-values and normalized enrichment score (NES). These were combined into a unified drug-level score by computing a concordance metric defined as:

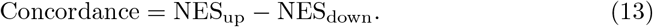

where NES_up_ corresponds to the normalized enrichment score obtained from the upregulated disease genes and NES_down_ corresponds to the enrichment score obtained from the downregulated genes. This formulation was chosen because effective therapeutic reversal requires simultaneously counteracting both components of the disease signature. A compound should ideally suppress genes that are upregulated in disease while restoring genes that are downregulated. Considering only a single enrichment score or selecting compounds solely based on the highest NES may fail to capture this bidirectional reversal effect. In particular, some compounds may strongly reverse one component of the signature while showing weak or even concordant behavior for the opposite component, resulting in biologically less meaningful prioritization despite significant statistical enrichment. By explicitly combining both enrichment directions into a single concordance measure, the ranking better reflects the overall capacity of a compound to reverse the disease-associated transcriptional state.

To integrate evidence from both upregulated and downregulated signatures, p-values were combined using Fisher’s method:

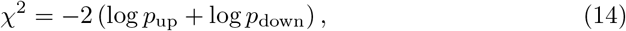

yielding a joint ranking statistic.

Drugs were ranked per sample based on the combined p-value (ascending) and concordance score (descending), yielding a prioritized list of candidate compounds for each transcriptional signature.

To ensure comparability with RANKOR, the candidate drug space was restricted to compounds present in the LINCS dataset with valid chemical structure annotations (SMILES), and overlapping with the evaluation set.

Importantly, this evaluation setup may lead to optimistic performance estimates for enrichment-based methods. Because the LINCS database serves both as the source of perturbational signatures and as the reference gene set collection, some test signatures may correspond to, or closely resemble, profiles already present in the background dataset. This introduces a potential form of information leakage, whereby the method may effectively recover known perturbations rather than generalizing to unseen transcriptional patterns. Consequently, the reported performance of GSEA should be interpreted as an upper-bound estimate under these conditions.

### 3.10 Baseline: ASGARD drug repurposing analysis for single cell data

To benchmark RANKOR against a representative gene-signature–based drug repurposing approach for single cell data, we implemented the ASGARD (He et al. 2023) baseline using the authors’ reference workflow and R package. ASGARD was selected because it is specifically developed to prioritize therapeutic compounds from single-cell–derived gene signatures and has been shown to outperform classical connectivity-based approaches by incorporating a refined drug scoring scheme. In particular, ASGARD evaluates the negative connectivity between a query disease signature and reference drug perturbation profiles, while emphasizing concordance among consistently regulated genes.

We constructed an ASGARD reference from LINCS L1000 Level 5 z-score signatures exported from our LINCS matrix. The resulting matrix was converted to within-instance gene ranks (descending by z-score), producing the ASGARD `rankMatrix`. Using the ASGARD R package, we then created the drug reference object with `GetDrugRef` and stored it as a serialized R object for reuse across experiments.

For each dataset, we computed query signatures at the patient × cell-state resolution using pseudo-bulked expression profiles. In the T-ALL analysis, cells were grouped by patient and Leiden cluster, expression was averaged per cluster, and per-gene z-scores were computed relative to matched healthy reference cells within the same cluster, according to the RANKOR preprocessing strategy, to ensure comparability. In the glioblastoma analysis, cells were grouped by patient, treatment condition, and annotated cell type (with tumor cells further stratified by tumor state), and pseudo-bulk means were computed per group. Query signatures were restricted to the LINCS gene set and represented as signed gene scores.

#### ASGARD scoring and drug ranking

For each patient-specific cell state, the signed z-score vector was supplied to ASGARD as the query gene score and the connectivity was computed against the LINCS-derived reference. Drugs were then ranked by the ASGARD significance metric.

For analyses focusing on known perturbations (etoposide in glioblastoma and doxorubicin in T-ALL), we reported the rank of the target drug within each patient. All ASGARD analyses were executed using the provided Asgard software container (run with Singularity) to ensure reproducibility.

## 4 Discussion and Conclusion

RANKOR is designed to take a disease-associated transcriptomic signature and return a ranked list of candidate therapies. The broader vision was to provide a machine learning framework that can support transcriptomics-guided therapeutic prioritization across bulk and single-cell settings, while remaining sufficiently interpretable and flexible for translational use.

The main contribution of this study is the formulation of drug prioritization itself as the primary modeling objective. To our knowledge, RANKOR is among the first machine learning frameworks explicitly designed to address the transcriptomics-based signature reversal problem directly, rather than indirectly through prediction of high-dimensional gene expression profiles, potency metrics, or post hoc similarity analysis. This distinction is important. In clinical and translational practice, the relevant question is often not how a drug perturbs a cell, but which drugs should be prioritized for a disease-associated molecular state.

Across the bulk perturbational benchmarks, RANKOR showed strong and consistent performance. In some settings it outperformed the standard enrichment-based baseline GSEA, whereas in others it achieved broadly comparable performance. We consider this an encouraging result rather than a replacement claim. GSEA remains a strong and highly interpretable standard for transcriptomic drug prioritization. However, we note that the ranking obtained with GSEApy are likely over-optimistic as we cannot exclude data leakage from test samples to the pre-defined drug consensus signatures. At the same time, the latent-space formulation of RANKOR appears advantageous when more detailed transcriptional structure is informative, particularly for distinguishing among individual compounds that share partially overlapping pathway effects.

However, a unique advantage of RANKOR is its ability to prioritize transcriptionally unseen drugs. Classical connectivity-based and enrichment-based approaches fundamentally depend on the availability of reference perturbational signatures for each compound. In contrast, RANKOR can embed a compound directly from its chemical structure and include it in the ranked candidate set even if no transcriptomic profile has previously been measured for that drug. This capability is particularly relevant for repurposing and translational discovery, where many potentially interesting compounds are not represented in existing perturbational atlases.

The explainability analyses further support the biological plausibility of the framework. This shows that the model is not relying on arbitrary transcriptomic artifacts, but rather on coordinated transcriptional programs linked to drug targets and downstream pathway responses. Such interpretability is important for trust and adoption, especially in settings where ranked therapeutic hypotheses may guide further laboratory or clinical evaluation.

At the same time, the results also highlight clear limitations. First, RANKOR depends on the presence of meaningful, drug-specific transcriptional signal. When perturbations induce only weak targeted changes, or when broad secondary and stress-related effects dominate the expression profile, the model has limited information from which to infer the correct therapeutic mapping. This was evident in the dose-dependent analyses, where prioritization improved substantially as the transcriptional response became stronger and more mechanistically informative. In that sense, RANKOR does not overcome the biological limits of the input data; rather, its performance remains bounded by signal quality and perturbation specificity.

Second, the chemical space used here was constructed from LINCS-associated compounds using a hierarchical mechanism-aware strategy. While this design was effective for the present study, it remains a relatively compact and task-specific chemical representation. Richer molecular encoders trained on broader chemical corpora and additional molecular properties may improve discrimination between structurally or pharmacologically similar compounds. In future work, it would be interesting to replace or augment the current chemical encoder with more general molecular foundation models, including recent approaches such as CheMeleon that learn transferable molecular representations from large descriptor-based pretraining corpora (Burns et al. 2025).

Third, the current framework prioritizes monotherapies. In many translational settings—particularly in oncology—clinically useful treatment strategies rely on drug combinations, both to improve efficacy and to reduce toxicity through lower individual exposures. Modeling drug synergy remains outside the scope of the present framework, but it is a natural and important direction for future work (Pan et al. 2023; Duarte and Vale 2022). Extending RANKOR toward combination prioritization would substantially increase its clinical relevance.

A related limitation is that RANKOR prioritizes compounds but does not currently suggest clinically realistic dosing strategies. For translational deployment, ranked therapies would ideally be integrated with pharmacokinetic and pharmacodynamic modeling, so that molecular prioritization could be linked to feasible exposure ranges, dosing schedules, and toxicity constraints in the human body (Papachristos et al. 2023;Liao et al. 2021). Such integration could help bridge the gap between transcriptomic prioritization and practical therapeutic decision-making.

Another aspect to be considered is that failure to correctly prioritize unseen drugs is not always attributable to meaningful biological differences. In some cases, it may arise from the presence of pharmacologically similar compounds with overlapping chemical structures or shared mechanisms of action, making precise discrimination challenging. In other instances, it may reflect limitations in the training data, where certain mechanisms are insufficiently represented across cellular contexts, leading to incomplete learning of related drug-specific transcriptional patterns.

Finally, although the present results support the utility of RANKOR across bulk and single-cell data, broader validation will be needed across additional diseases, perturbational platforms, and patient-derived cohorts. In particular, its performance in settings with more subtle disease signals, stronger microenvironmental confounding, or limited reference coverage remains to be established. Future benchmarking against additional clinical and functional drug-screening datasets will be important for assessing generalizability and translational utility.

Overall, RANKOR should be viewed as a machine learning alternative for transcriptomics-based drug prioritization that is especially attractive when one seeks direct ranking, scalability, and extension to unseen compounds. Rather than replacing existing enrichment-based approaches, it provides a complementary strategy that appears particularly well suited to settings where transcriptomic magnitude, structure, and cross-modal generalization matter.

## Data and Code Availability

The data used in this study is described in the manuscript. The source code for data processing and analysis is publicly available at: RANKOR

